# The mGlu7 receptor provides protective effects against epileptogenesis and epileptic seizures

**DOI:** 10.1101/514844

**Authors:** Benoit Girard, Pola Tuduri, Maria Paula Moreno, Sophie Sakkaki, Cedric Barboux, Tristan Bouschet, Annie Varrault, Jihane Vitre, Isabelle McCort, Julien Dairou, Francine Acher, Laurent Fagni, Nicola Marchi, Julie Perroy, Federica Bertaso

## Abstract

Finding new targets to control or reduce seizure activity is essential to improve the management of epileptic patients. We hypothesized that activation of the pre-synaptic and inhibitory metabotropic glutamate receptor type 7 (mGlu7) reduces spontaneous seizures.

We tested LSP2-9166, a recently developed mGlu7/4 agonist with unprecedented potency on mGlu7 receptors, in two paradigms of epileptogenesis. In a model of chemically induced epileptogenesis (pentylenetetrazol systemic injection), LSP2-9166 induces an anti-epileptogenic effect rarely observed in preclinical studies. In particular, we found a bidirectional modulation of seizure progression by mGlu4 and mGlu7 receptors, the latter preventing kindling. In the intra-hippocampal injection of kainic acid mouse model that mimics the human mesial temporal lobe epilepsy, we found LSP2-9166 reduced seizure frequency and hippocampal sclerosis. LSP2-9166 also acts as an anti-epileptic drug on established seizures in both models tested.

Specific modulation of the mGlu7 receptor could represent a novel approach to reduce pathological network remodeling.

## INTRODUCTION

Epilepsy is a predisposition of neural networks to generate spontaneous and recurrent seizures (Fisher et al., 2014). Epileptogenesis defines the ensemble of processes that, starting from a given genetic environment and/or precipitating factors, lead to the formation of a neuronal network supporting hyperexcitability and hypersynchronization (Terrone et al., 2016). Current antiepileptic drugs (AED) allow seizure control only in 70% of patients (Asadi-Pooya et al., 2017; Picot et al., 2008), most have debilitating side effects and they do not necessarily treat the co-morbidities related to the pathology (e.g. anxiety, depression, cognitive deficits) (Chen et al., 2017). Furthermore, treating the underlying mechanisms to achieve the prevention or remission of epileptogenesis remains a challenge (Terrone et al., 2016). The research of new targets in epilepsy is therefore crucial if we want to outweigh the current limits of classical AED families.

The metabotropic glutamate receptor type 7 (mGlu7) is widely expressed in key regions of the central nervous system involved in different epilepsies syndromes including the thalamocortical loop, the hippocampus or the amygdala (Bradley et al., 1998). The mGlu7 receptor is located in the active zone of glutamatergic and GABAergic presynapses (Kinoshita et al., 1998; Somogyi et al., 2003). Its activation induces a decrease of neurotransmitter release through inhibition of calcium currents (Millán et al., 2002; Perroy et al., 2000).

The pharmacology of group III mGluR (mGlu4, 6, 7 and 8) has slowly evolved, due to the strong homology with other members of the metabotropic glutamate receptor family (Pin and Duvoisin, 1995). The use of transgenic animals has highlighted a dynamic control of synaptic transmission and plasticity by mGlu7. For example, in the hippocampus CA1 area or the amygdala, mGlu7 is necessary for the disinhibition of GABAergic neurons allowing the induction of LTP between SC-CA1 synapses (Klar et al., 2015) or thalamic input, respectively (Fendt et al., 2013). Constitutive mGlu7 receptor activity at specific thalamic synapses avoids pathological recurrent oscillatory network activity (Tassin et al., 2016). In mGlu7 KO mice, the absence of the receptor induces an aberrant increase in the theta power rhythms during spatial memorization tasks preventing effective integration of information (Holscher et al., 2005). Spontaneous tonico-clonic epileptic seizures have also been observed in these mice (Sansig et al., 2001). The receptor mutation in mGlu7^AAA^ KI mice disrupting a PICK1-related signaling pathway leads to spontaneous absence-like seizures (Bertaso et al., 2008) and increases susceptibility to pro-convulsive drugs such as pentylentetrazole (PTZ) (Zhang et al., 2008). Overall, the results suggest that mGlu7 acts as a buffer to maintain relevant oscillation patterns avoiding hyperexcitability and hypersynchronization of neural networks.

We postulated that exogenous activation of the mGlu7 receptor in pathological conditions associated with epileptogenesis could reduce and possibly prevent the onset of epileptic seizures. To test this hypothesis we needed pharmacological tools that allow targeting mGlu7 receptors. Thanks to a virtual drug screening followed by hit optimization, we have developed new agonists with unprecedented potency for mGlu7 (Acher et al., 2012). These compounds are water-soluble, they cross the blood-brain barrier, show a broad selectivity (Belhocine et al., 2018) and have demonstrated efficacy in animal models of opiates/alcohol dependence and pain (Hajasova et al., 2018; Lebourgeois et al., 2018; Vilar et al., 2013).

We chose to investigate the effect of LSP2-9166 on two complementary models of epilepsy in mice. The chemical kindling induced by pentylenetetrazol (PTZ), a non competitive GABAA receptor antagonist, is a recognized model of primary generalized tonico-clonic seizures (Kupferberg, 2001). Kindling corresponds to a progressive sensitization to pro-epileptic stimulation. The model consists of repeated injections of a sub-convusive dose of PTZ that induces a progressive intensification of epileptic activity, from absence seizures to generalized tonic-clonic seizures. The advantage of this model is undoubtely the temporal control of seizure occurrence and the highly standardized scoring system. A previous study showed that the non-specific group III mGlu agonist, L-AP4, has no effect on PTZ kindling in rats (Maciejak et al., 2009). However, the concentrations used did not guarantee the activation of the low-affinity mGlu7 receptor, whose EC50 for L-AP4 is around 200 μM (Selvam et al., 2018).

Temporal lobe epilepsy (TLE) is the most common form of complex partial epilepsy affecting 60% of epileptic patients. The patients show hippocampal sclerosis characterized by neuronal loss in the CA1/CA3 region of the hippocampus and hilus with granular cells dispersion and mossy fibers sprouting in the molecular layer and the dentate gyrus (Asadi-Pooya et al., 2017; Tellez-Zenteno and Hernandez-Ronquillo, 2012). Intra-hippocampal injection of kainic acid in rodents provides a model which effectively recapitulates most of the clinical features of TLE (Bouilleret et al., 1999), including pharmacoresistance. These two models allowed us to study not only the impact of mGlu7 activation on seizures but also on the development of the pathology and the associated behavioural alterations. A reduction in both chemical kindling progression and TLE seizures was observed upon acute or repeated administration of the agonist.

## RESULTS

### Chronic administration of LSP2-9166 has no negative side effects on mice behaviour

Before setting out to study the activation of the mGlu7 receptor in epilepsy, we determined the adequate dose to be administered *in vivo*. As previously reported, LSP2-9166 (cited in the illustrations as “LSP2” for simplicity) has a potency (EC_50_) for mGlu7 receptors of 2.21±0.53 µM and 0.07±0.01 µM for mGlu4 receptors, as we tested in HEK cells overexpressing either receptors. Intraperitoneal injection of two different doses of LSP2-9166 showed a plasmatic concentration reaching its maximum 30 min after administration and returning to nearly undetectable levels within 2 h (Fig. 1A). In addition, based on a compound belonging to the same chemical family (LSP1-2111, (Cajina et al., 2014) and on the pharmacodynamics features of LSP2-9166, reasoned that 1 mg/kg of LSP2-9166 should predominantly activate mGlu4 receptors whereas 10 mg/kg would reach the EC_50_ for mGlu7. These parameters provided an operational window for the acute effects of the compound.

**Figure 1:**
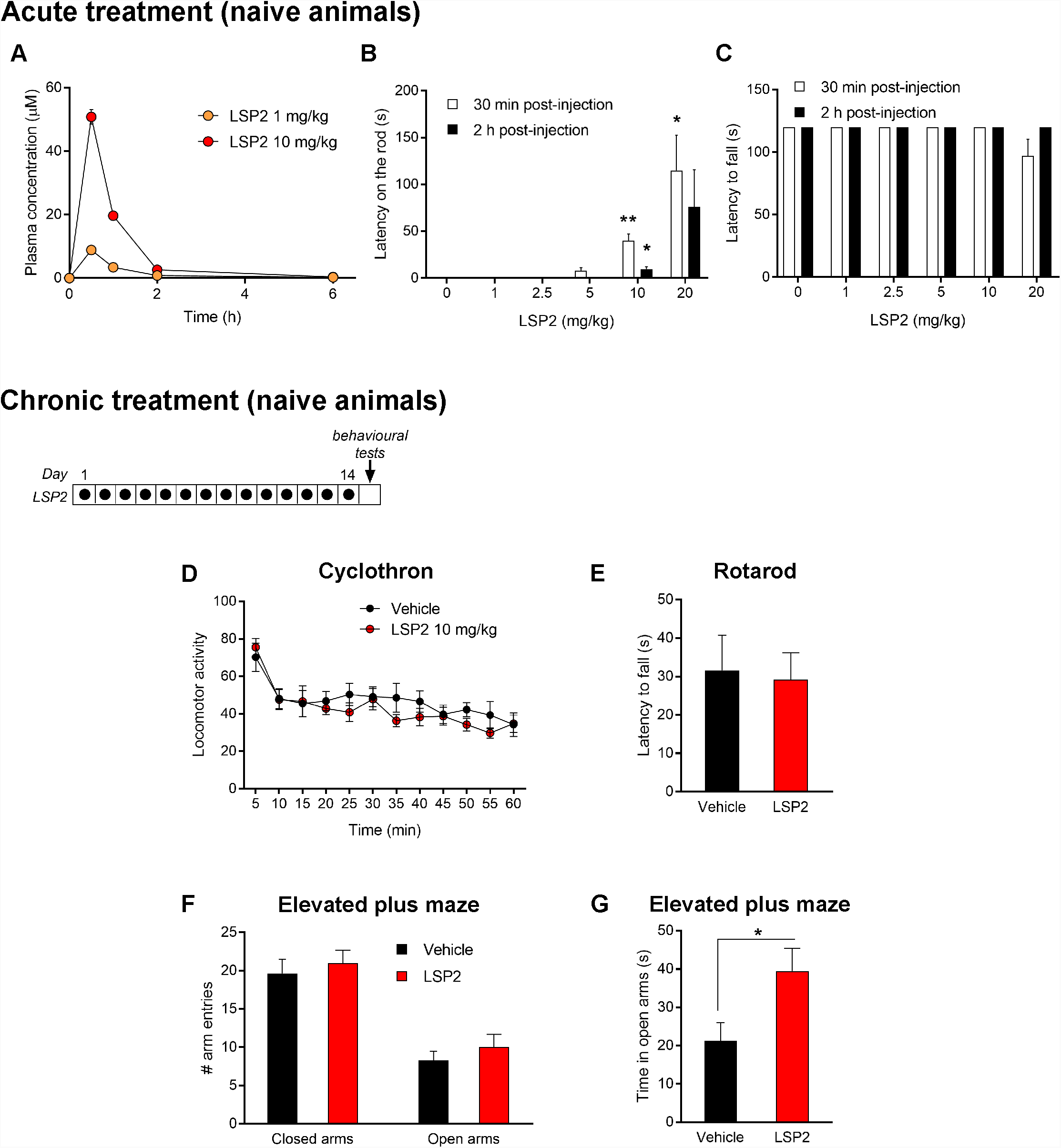
Acute and chronic administration of LSP2-9166 on naive animals. (A) Pharmacokinetics of two different doses of LSP2-9166 (LSP2 for simplicity hereafter) in the plasma (n = 4 per time point and dose). (B) Latency to move from the rod 30 min and 2 h after acute injection of different doses of LSP2 (n = 4 to 8). (C) Latency to fall off the rotarod 30 min and 2 h after acute injection of different doses of LSP2 (n = 4 to 8). (D-G) Behavioural effects on naïve animals after a 2-week daily treatment with LSP2 10 mg/kg (n = 8 for all conditions and tests). (D) Locomotor activity measured over 1 h in the circular corridor. (E) Latency to fall from the rotarod. (F, G) Elevated plus maze activity as number of entries in closed and open arms and time spent in each type of arm. *, p< 0.05; **, p< 0.01, Mann-Whitney test.

We first tested the effects of acute systemic administration of LSP2-9166. At the dose of 10 mg/kg the compound was associated to a small transient inactive state resembling catalepsy, as showed by the slower response of the treated animals when their forepaws where set on a rod compared to controls (Fig. 1B). However, mice were readily responding to sound visual or touch stimulation, and motor coordination was not affected, as indicated by the rotarod motor performance values (Fig. 1C).

AED are typically administered on a daily base and their side effects are in most cases problematic for the patients’ quality of life. We tested the effect of daily systemic administration of LSP2-9166 (10 mg/kg i.p.) on naive animals over 14 days. No difference was detected in any of the paradigms used, including the circular corridor for locomotor activity (Fig. 1D), or rotarod performance test (Fig. 1E). Anxiety levels were higher in the time spent in the open arms with no change in the number of entries in either type of arm (Fig. 1F, G), reflecting a possible anxiolytic effect. The levels of *Grm7* or *Grm4* mRNAs in either the cortex or the hippocampus were unaffected by systemic administration of LSP2-9166 (Table 2).

**Table 2.** RT-QPCR quantification. Quantification of different gene expression in naive animals after a 2-week treatment with LSP2-9166 (10 mg/kg) or vehicle; in mice after the PZ kindling protocol, with or without daily treatment with LSP2-9166 (10 mg/kg); in IHKA mice at 14 or 28 days post-KA injection, with or without bi-daily treatment with LSP2-9166 (10 mg/kg) during 14 days. Where no value of p is indicated, p > 0.05 (Mann-Whitney test). n = 4 for all conditions.

| Gene | Region |  |  |  |  |  |
| --- | --- | --- | --- | --- | --- | --- |
| **Chronic treatment (naive animals)** |  | **Vehicle** | **LSP2-9166** |  |  |  |
| *Grm7* | Cortex | $0.653 \pm 0.041$ | $0.699 \pm 0.023$ | | | |
| | Hippocampus | $0.495 \pm 0.046$ | $0.529 \pm 0.023$ | | | |
| *Grm4* | Cortex | $0.185 \pm 0.014$ | $0.225 \pm 0.004$ (* $p = 0.027$) | | | |
| | Hippocampus | $0.042 \pm 0.004$ | $0.036 \pm 0.003$ | | | |
| *Fabp7* | Cortex | $0.188 \pm 0.018$ | $0.181 \pm 0.009$ | | | |
| | Hippocampus | $0.113 \pm 0.008$ | $0.097 \pm 0.010$ | | | |
| *mKi67* | Cortex | $0.006 \pm 0.001$ | $0.005 \pm 0.001$ | | | |
| | Hippocampus | $0.005 \pm 0.0003$ | $0.005 \pm 0.001$ | | | |
| *Pdgfrb* | Cortex | $0.200 \pm 0.012$ | $0.183 \pm 0.017$ | | | |
| | Hippocampus | $0.150 \pm 0.009$ | $0.153 \pm 0.009$ | | | |
| **PTZ kindling model** |  | **Naive** | **PTZ** | **PTZ + LSP2-9166** |  |  |
| *Grm7* | Cortex | $1.522 \pm 0.670$ | $1.795 \pm 0.310$ | $1.518 \pm 0.303$ | | |
| *Grm4* | Cortex | $0.131 \pm 0.023$ | $0.199 \pm 0.028$ | $0.346 \pm 0.090$ | | |
| *Fabp7* | Cortex | $0.235 \pm 0.088$ | $0.364 \pm 0.130$ | $0.166 \pm 0.050$ | | |
| *mKi67* | Cortex | $0.022 \pm 0.012$ | $0.028 \pm 0.003$<br>(** $p = 0.003$ vs. Naive) | $0.030 \pm 0.009$<br>(* $p = 0.04$ vs. PTZ) | | |
| *Pdgfrb* | Cortex | $0.167 \pm 0.028$ | $0.125 \pm 0.008$ | $0.168 \pm 0.012$<br>(* $p = 0.03$ vs. PTZ) | | |

| IHKA model |  | Naive | IHKA D14 Vehicle | IHKA D14 LSP2-9166 | IHKA D28 Vehicle | IHKA D28 LSP2-9166 |
| --- | --- | --- | --- | --- | --- | --- |
| *Grm7* | Hipp Ipsi | 0.659 ± 0.052 | 1.192 ± 0.196<br>(** p = 0.006 vs. Naive) | 1.327 ± 0.077 | 1.576 ± 0.588<br>(** p = 0.007 vs. Naive) | 0.991 ± 0.431 |
|  | Hipp Contra | 0.650 ± 0.026 | 0.774 ± 0.090 | 0.699 ± 0.079 | 1.386 ± 0.208<br>(*** p = 0.0002 vs. Naive) | 1.323 ± 0.776 |
| *Grm4* | Hipp Ipsi | 0.085 ± 0.010 | 0.178 ± 0.032<br>(** p = 0.006 vs. Naive) | 0.165 ± 0.019 | 0.146 ± 0.035<br>(* p = 0.04 vs. Naive) | 0.156 ± 0.041 |
|  | Hipp Contra | 0.059 ± 0.007 | 0.098 ± 0.021<br>(* p = 0.04 vs. Naive) | 0.098 ± 0.025 | 0.076 ± 0.022 | 0.096 ± 0.033 |
| *Fabp7* | Hipp Ipsi | 0.227 ± 0.038 | 0.303 ± 0.041 | 0.239 ± 0.077 | 0.404 ± 0.083 | 0.328 ± 0.057 |
|  | Hipp Contra | 0.205 ± 0.044 | 0.158 ± 0.007 | 0.699 ± 0.079 | 0.390 ± 0.208 | 0.264 ± 0.024 |
| *mKi67* | Hipp Ipsi | 0.010 ± 0.002 | 0.010 ± 0.002<br>(** p = 0.005 vs. Naive) | 0.022 ± 0.004<br>(* p = 0.013 vs. Naive) | 0.013 ± 0.006 | 0.014 ± 0.005 |
|  | Hipp Contra | 0.011 ± 0.001 | 0.018 ± 0.003<br>(* p = 0.03 vs. Naive) | 0.012 ± 0.002 | 0.018 ± 0.007 | 0.022 ± 0.010 |
| *Pdgfrb* | Hipp Ipsi | 0.135 ± 0.014 | 0.238 ± 0.008<br>(*** p = 0.0006 vs. Naive) | 0.185 ± 0.017<br>(* p = 0.03 vs. D14 Vehicle) | 0.052 ± 0.017<br>(* p = 0.010 vs. Naive) | 0.159 ± 0.035 |
|  | Hipp Contra | 0.114 ± 0.020 | 0.149 ± 0.015 | 0.141 ± 0.016 | 0.074 ± 0.017 | 0.088 ± 0.003 |

### LSP2-9166 neutralizes chemical kindling

The effects of systemic injection of LSP2-9166 on epileptogenesis were first determined during chemical kindling. We determined the subthreshold dose of pentylenetetrazol (PTZ) to be used by looking at the response of mice to a single dose of PTZ (30 to 60 mg/kg i.p.). The chemical convulsant produces clear-cut EEG and behavioural signatures (Fig. 2A) that can be readily used to score the animal reactions. Mice displayed partial clonic seizures for doses greater than 40 mg/kg; this dose was therefore chosen as subthreshold to start the kindling protocol. We then asked whether LSP2-9166 could have an acute effect on PTZ administration, independent of the processes underlying epileptogenesis. Pre-treatment with LSP2-9166 (10 mg/kg i.p. 20 min prior to PTZ injection) produced no difference in the mean score and latency between vehicle- and LSP2-9166-treated animals, whatever the dose of PTZ tested (Fig. 2B, C), with only a delayed latency of the response to PTZ observed at 60 mg/kg (Fig. 2C). This result suggested that the kindling would not be affected by a direct block of PTZ-induced seizures.

**Figure 2:**
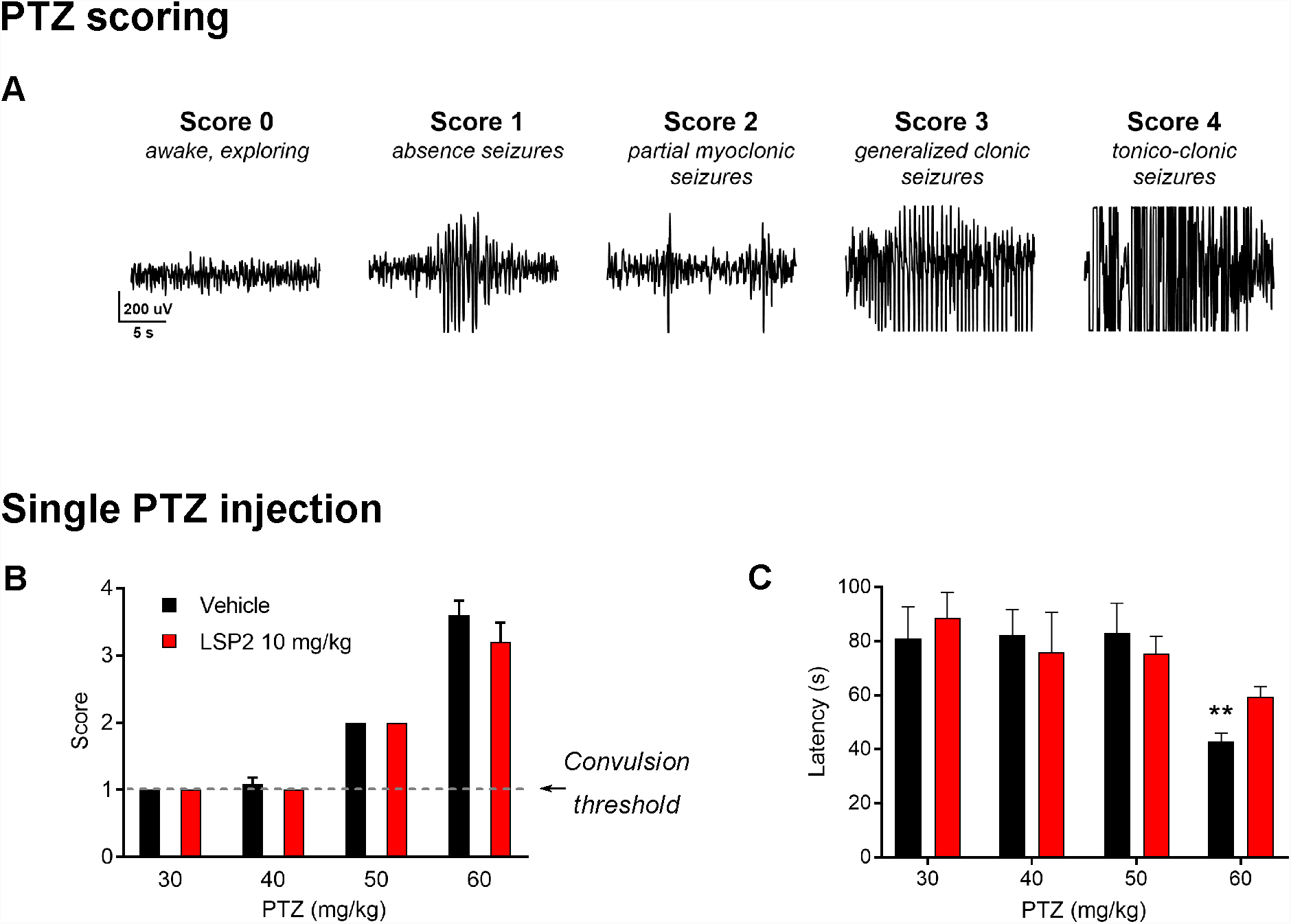
Acute effect of LSP2 on simple PTZ-induced seizures. (A) Example of EEG recordings representative of the different severity scores with a description of the main phenotype observed. (B, C) Effect of LSP2 pre-treatment on severity score (B) and latency to develop such score (n = 3 to 10 per condition). (C) of seizures induced by different acute doses of PTZ. **, p< 0.01, Mann-Whitney test.

Kindled animals received a daily injection of LSP2-9166 (1 or 10 mg/kg, i.p.) or vehicle (PBS). From day 1 and every other day, anials received a the sub-convulsive dose of PTZ (40 mg/kg, i.p.), their behaviour was observed and their EEG activity recorded. Control animals injected with PBS underwent the expected gradual, long-lasting seizure development whose severity increased from simple prostration on day 1 to tonico-clonic seizures on day 15 (Fig. 3A). Treatment with the higher dose of LSP2-9166 (10 mg/kg) completely blocked the progression of kindling, whereas the lower dose (1 mg/kg) had no significant effect. Analysis of scores distribution between day 13 and 15 of kindling shows that in both control and low-dose LSP2-9166-treated groups, 75% of the mice experienced convulsive seizures corresponding to at least score 2 or higher (Fig. 3B). Treatment with LSP2-9166 (10 mg/kg) inversed the distribution: only 15% of the animals reached score 2 and 1.5% score 3. Thus, the lowest dose of LSP2-9166 (1 mg/kg) had no significant effect while with the highest LSP2 dose (10 mg/kg) completely blocked the progression of kindling. This protective effect lasted beyond the end of the treatment: a PTZ challenge 30 days after the last PTZ injection showed that kindling kept progressing in the control group, while the LSP2-9166-treated group still produced a low response, without further injection of LSP2-9166.

**Figure 3:**
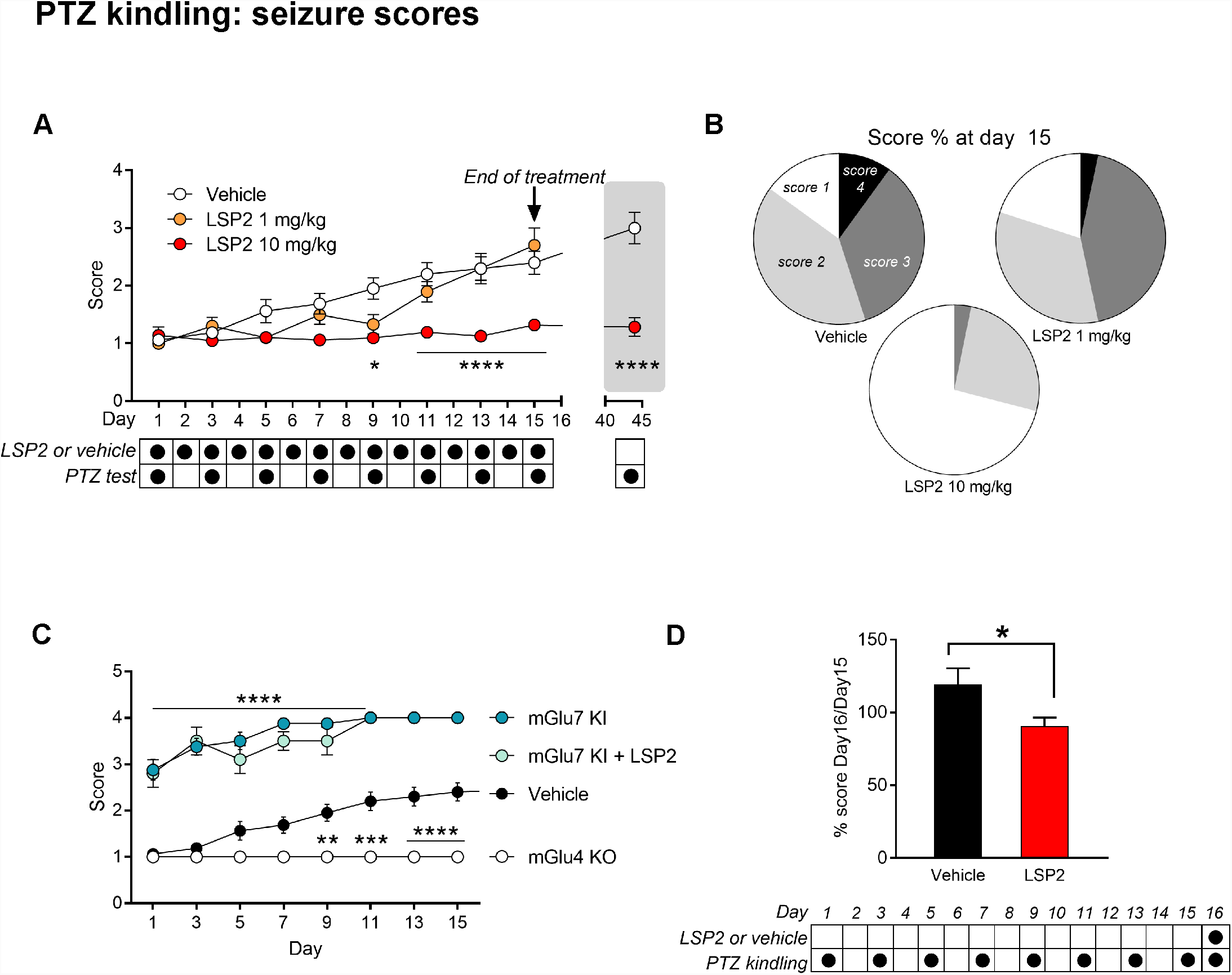
Chemical PTZ kindling is attenuated by mGlu7 activation. (A) Progression of the severity score during PTZ kindling upon the indicated treatments. LSP2 or vehicle were administered daily while the PTZ subthreshold dose was administered every two days. (B) Distribution of the score severity depending on the dose of LSP2 administered. (C) Progression of the severity score during PTZ kindling in different mouse genotypes (n = 7 to 16). *, p< 0.05; **, p< 0.01, ***, p< 0.001, ****, p< 0.0001, two-way ANOVA. (D) Effect of LSP2 injection on the evolution of kindled seizure induced by an acute subconvulsive dose of PTZ at the end of the kindling. *, p< 0.05, Mann-Whitney test.

We have previously shown that mGlu7^AAA^KI mice (mGlu7 KI in the figures for simplicity) show an increased susceptibility to PTZ, responding with convulsive seizures to a single, sub-threshold dose of the compound (40 mg/kg i.p.) (Zhang et al., 2008). Induction of PTZ kindling in mGlu7^AAA^ KI mice has devastating effects with rapid progression towards tonico-clonic seizures often followed by death (70% after the second administration of PTZ) (Fig. 3C). Treatment of mGlu7^AAA^ KI mice with LSP2-9166 had no protective effect on the kindling. On the other hand, knock-out of the other LSP2-9166 main target, the mGlu4 receptor, results in complete resistance to PTZ kindling with the totality of mGlu4 KO mice showing only prostration and absence-like EEG activity for the whole kindling protocol. These results point at a diametrically opposite effect of the two targets of LSP2-9166 on PTZ kindling.

We then asked whether LSP2-9166 is effective on kindled-seizures, i.e. does a single dose of LSP2-9166 reduce the response to PTZ in animals already sensitized to the convulsant? While a control injection showed the continuous progression of the score at day 16 vs. day 15, a single injection of LSP2-9166 reduced the score (Fig. 3D). This result shows that LSP2-9166 shapes primarily generalized seizures brought by repeated PTZ administration during their emergence and once instated.

During the kindling process, astrocytes can be activated independently of neuronal degeneration (Khurgel et al., 1995). Other epilepsy models have shown the occurrence of radial glia scaffolds favoring neuronal migration. An increase of the conversion of neural stem cells into reactive astrocytes has been reported in certain models of neuronal hyperactivation (Sierra et al., 2015). On the other hand, epileptogenesis is accompanied by neuroinflammation related to microglia activation and proliferation. Furthermore, the integrity of the blood-brain barrier can be compromised in certain types of epilepsy where it can have a causal role in seizure occurrence (Milesi et al., 2014). Immunohistochemical analysis indicate that treatment with LSP2-9166 at 10 mg/kg significantly reduced microglia cell density in the CA1 and CA3 hippocampal regions of kindled mice (IBA1, Fig. 4A, B). Astrocytosis, measured by GFAP staining intensity, was decreased in CA3 and DG (Fig. 4 C, D). Other markers related to epileptogenesis were assessed by quantitative RT-PCR (Table 2). The mRNA for fatty acid-binding protein 7 (*Fabp7*), a marker for radial glia, showed only a tendency to an increase in kindled animals, whereas the proliferation marker *mKi67* and the pericytic marker *Pdgfrb* (Milesi et al., 2014) mRNA are significantly decreased. All these changes were reversed by LSP2-9166 treatment.

**Figure 4:**
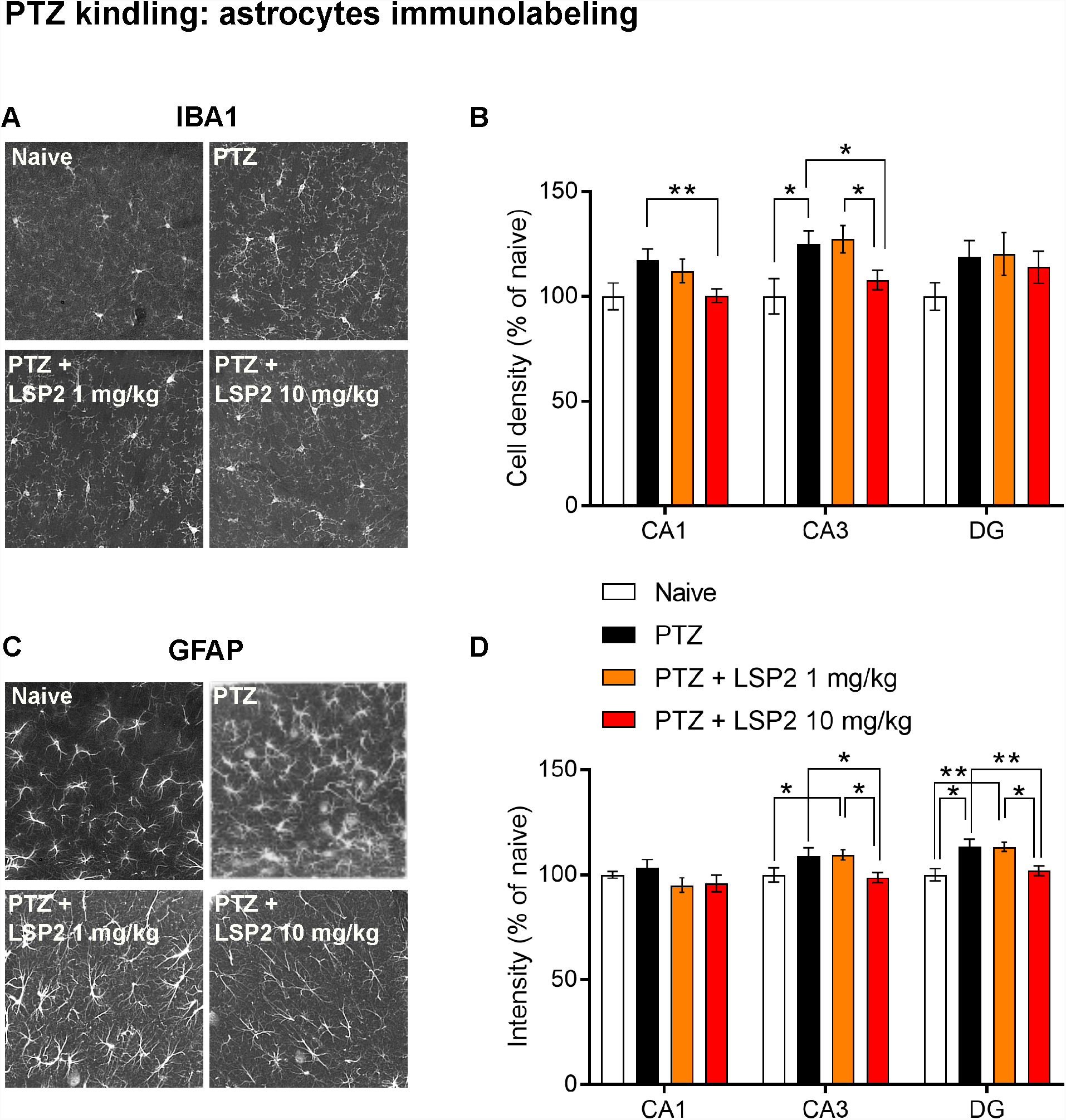
Astrocytosis and microglial activation in PTZ kindling. (A) Example of images of anti-IBA1 labelling representing different treatment effect on the microglial cell density. (B) Modification of microglial cell density in different hippocampal regions depending on the indicated conditions. (C) Example of images of anti-GFAP labelling representing different treatment effect on astrocyte activation. (D) Modification of astrocyte activation in different hippocampal regions depending on the indicated conditions. (n = 5 to 16 mice per condition, n = 2-4 slices per mouse). *, p< 0.05; **, p< 0.01, Mann-Whitney test.

Behavioural tests were performed before and after the kindling protocol. PTZ kindling reduced the locomotor activity in both the circular corridor test (Fig. 5A, B) and elevate plus maze, the latter showing a strong decrease in the number of visits regardless of the type of arm (open or closed, Fig. 5C). No difference was observed on the rotarod performance test, which avoids any passive behavior (Fig. 5D). Treatment with the LSP2-9166 had no significant effect on any of these tests, despite a non-significant increase of locomotion towards pre-kindling values. Interictal brain activity, measured one day after the last PTZ injection (day 16), showed that basal EEG spectrum of PTZ-kindled mice is modified, with an increase in the gamma band (Fig. 5E). LSP2-9166 treatment showed a tendency to normalize the interictal spectrum. For a target to be suitable for therapy, it needs to be available even in pathological conditions. *Grm7* and *Grm4* mRNAs were unaffected by all treatments, as suitable (Table 2).

**Figure 5:**
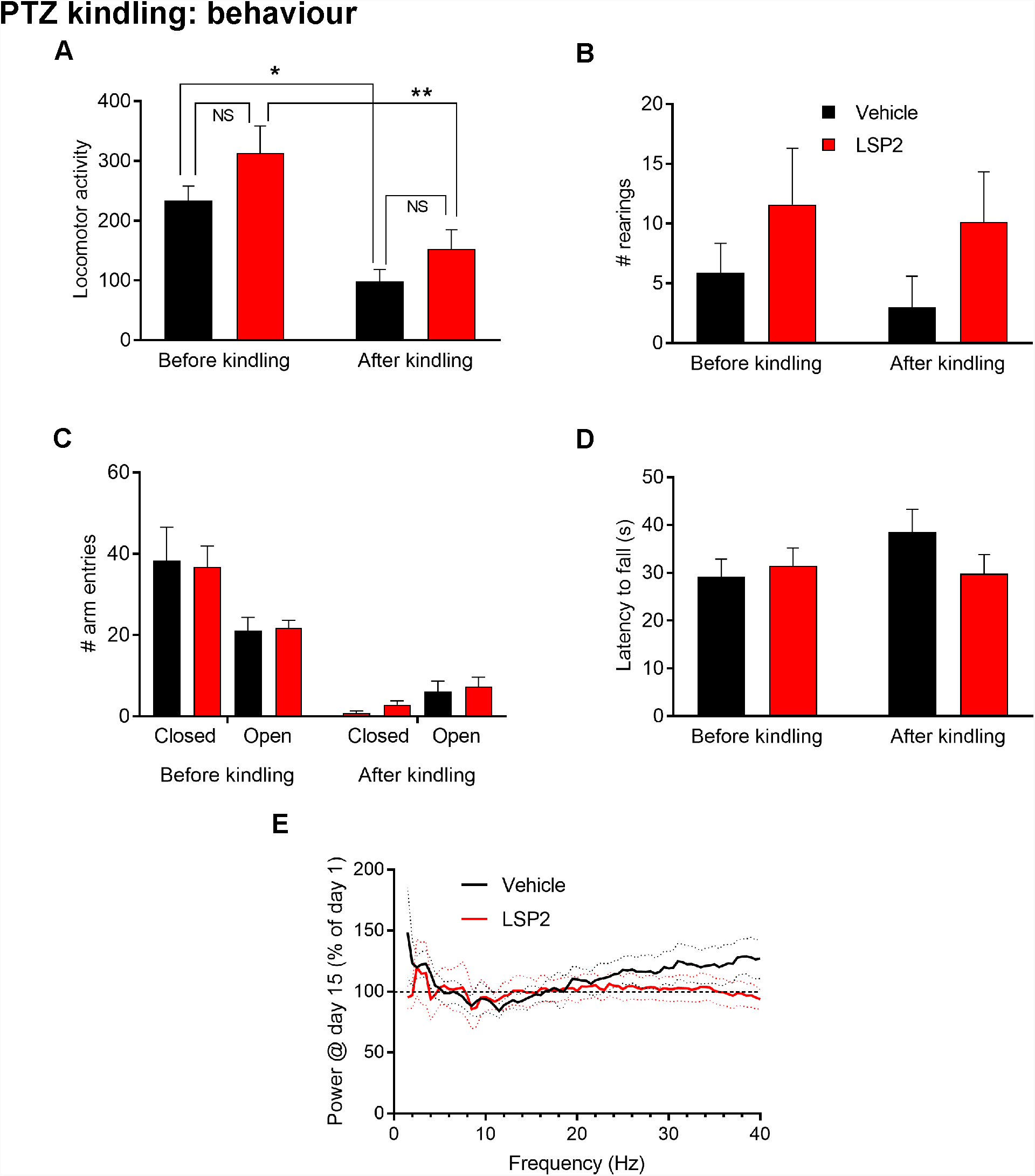
Behavioural effects of LSP2-9166 on PTZ-kindled behaviour. (A, B) Effect of LSP2 treatment on the evolution of locomotor activity (A) and rearings (B) before and after PTZ kindling. (C) Number of open and closed arm entries in the elevated plus maze. (D) Performance on the rotarod. (E) Effect of the LSP2-9166 treatment on the modification of the power spectrum induced by PTZ kindling. (n = 8 per condition and test). *, p< 0.05; **, p< 0.01, Mann-Whitney test.

### LSP2-9166 reduces seizures in the early phases of TLE progression

We then established the intra-hippocampal kainate (IHKA) injection model of mesial temporal lobe epilepsy in mice (Bouilleret et al., 1999). The animals were treated twice daily with either LSP2-9166 (10 mg/kg) or vehicle (PBS solution) during 15 days (Fig. 6A). Analysis of several parameters was performed for at least 5 weeks post-SE in order to characterize seizure progression. Visual observation of EEG recordings indicated that paroxysmal activity was detected as early as day 5, with an amplitude increasing swiftly over the following days. Overall, LSP2-9166 had an effect during the administration phase, where it significantly reduced the total duration of seizures and the probability to develop long seizures (more than 15 s) with a steady tendency for a smaller number of seizures (Fig. 6B, C). Suspension of the treatment caused a transient rebound in total seizure duration. Interestingly LSP2-9166 seems to generate a temporal spacing of seizures, as shown by the probability of the inter-seizure interval to last longer than 30 s, even after discontinuation of the treatment.

**Figure 6:**
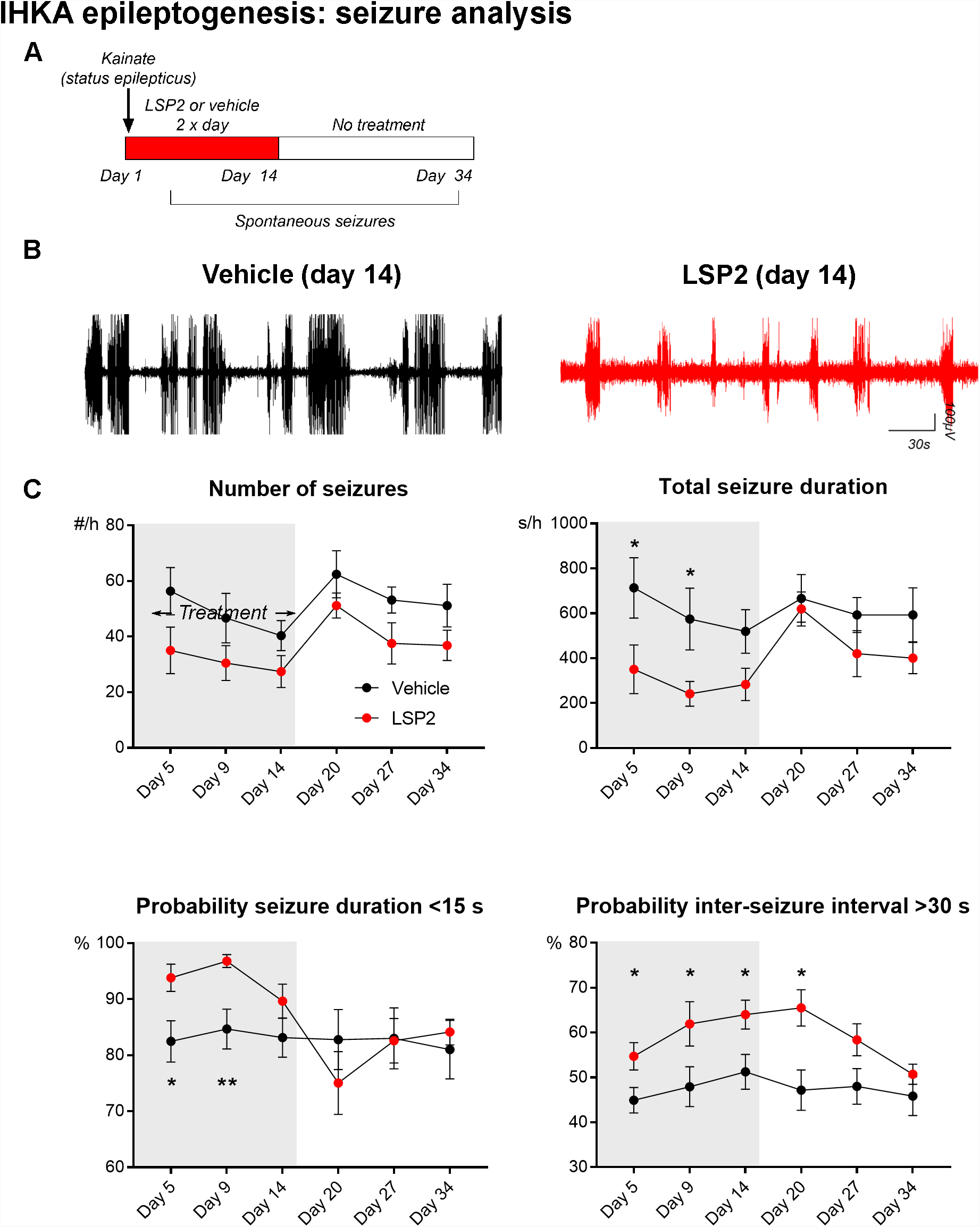
Transient effect of LSP2-9166 on IHKA-induced epileptogenesis. (A) Experimental protocol. (B) Example EEG recordings at day 14 from a vehicle-treated and an LSP2-treated mouse. (C) Effect of LSP2 treatment on the evolution of number of seizures per hour, total seizure duration per hour, probability of seizure duration shorter than 15 s, and probability of inter-seizure interval being shorter than 30 s. (n = 6 to 18 per condition). *, p< 0.05, **, p< 0.01, two-way ANOVA.

The IHKA model triggers an extensive damage of the hippocampus ipsilateral to the injection side (Fig. 7), especially marked by the dispersion of the granule cells of the dentate gyrus. As for the kindling model, we compared different markers of cell reactivity. During the early phase of epileptogenesis, a moderate astrocytosis was observed in the contralateral side of the hippocampus, more significantly in the CA1 area. LSP2-9166 reversed this phenomenon, whereas at day 28 it seemed to have an opposite effect (GFAP, Fig. 7C, D). The ipsilateral side showed an astrocytic activation only in the later phase, after LSP2-9166 treatment arrest (Fig. 7E). Microglial activation was striking on both ipsi- and contralateral sides of the hippocampus (Fig. 7B, G, H). LSP2-9166 treatment increased this trend in the contralateral side while inducing a significant decrease in the ipsilateral CA1 area. Iba1 staining decreased in the control group at day 28 and was not affected by LSP2-9166. The thickness of dentate gyrus doubled in both control and treated animals. However, while the dispersion continued reaching a 6-fold increased at day 28, it was strongly reduced by LSP2-9166 (Fig. 7F). The amount of both mGlu7 and mGlu4 mRNAs was increased (Table 2), as reported in human TLE (Lie et al., 2000), as was the proliferation marker *mki67*. Radial glia’s marker *Fabp7* showed a tendency to increase especially in the contralateral side, while changes in *Pdgfrb* might indicate disruption of BBB homeostasis in the KA-injected side (Table 2). No difference on overall behavioural performance including locomotor activity and motor coordination was induced by the treatment (Fig. 8).

**Figure 7:**
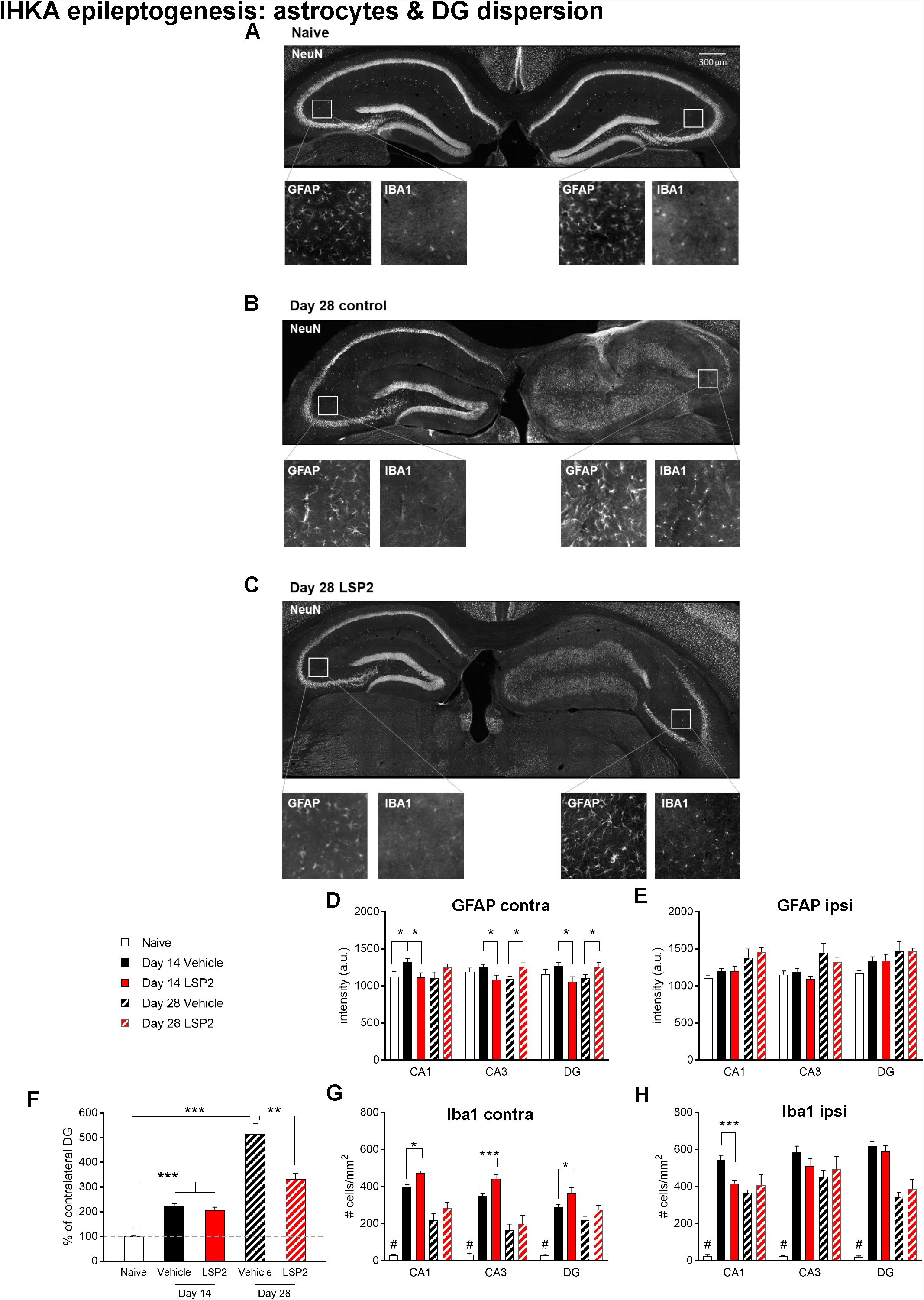
Immunohistochemical characterization of the late phase of IHKA epileptogenesis. (A-C) Example of images. For each panel, main image: hippocampus left, contralateral / hippocampus right, ipsilateral with anti-NeuN labelling; insets left to right, anti-GFAP contralateral, anti-IBA1 controlateral, anti-GFAP ipsilateral, anti-IBA1 ipsilateral. (A) Naive animal. (B) Control animal 28 days post-SE. (C) LSP2-treated animal 28 days post-SE. (D-E) Evolution of GFAP intensity post-SE in different regions of (D) contralateral and (E) ipsilateral hippocampus. (F) Effect of LSP2-9166 treatment on the evolution of dentate gyrus dispersion post-SE. (G-H) Effect of LSP2-treatment on the evolution of microglia cell density post-SE in different regions of (G) contralateral and (H) ipsilateral hippocampus. (n = 2 to 4 mice per condition, n = 4 slices per mouse). *, p< 0.05; **, p< 0.01, ***, p< 0.001, Mann-Whitney test.

**Figure 8:**
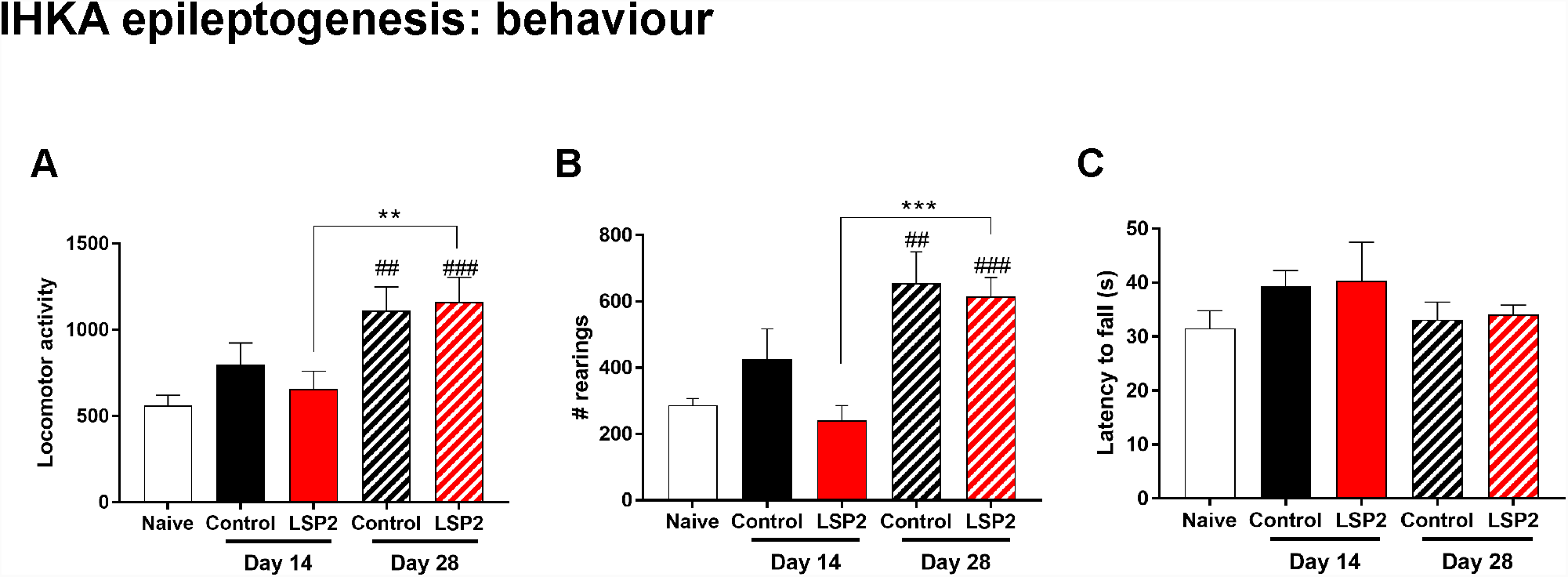
Behavioural effects of LSP2-9166 on IHKA-induced epileptogenesis. (A) Effect of LSP2 treatment on the evolution of locomotor activity induced 14 and 28 days post-SE. (B) Effect of LSP2-9166 treatment on the evolution of rearings. (C) Effect of LSP2 treatment on the performance on the rotarod. **, p< 0.01; ***, p< 0.001, Mann-Whitney test; ##, ###, different from naive condition.

### LSP2-9166 has an anti-epileptic effect on chronic hippocampal seizures

We asked whether LSP2-9166 could act as an anti-epileptic drug on established, chronic hippocampal seizures (Fig. 9A). Thirty days after SE, high (10 mg/kg), but not low (1 mg/kg, not shown) doses of LSP2-9166 significantly reduced the number and total duration of seizures (Fig. 9B, D, F, H), as well as the mean seizure duration, while it had no effect on peak amplitudes (Fig. 9I). The effect was comparable to that of the clinically used AED diazepam (DZP, 1 mg/kg), a benzodiazepine increasing GABAergic tone (Fig. 9C, D, F). However, the mechanism seems to be different. Firstly, the GABAA competitive antagonist flumazenil (FLU, 5 mg/kg) was able to significantly reduce the effect of DZP on seizure number and duration, but not that of LSP2-9166 (Fig. 9D, F). In addition, IHKA interictal activity showed a differential effect of the two compounds, DZP and LSP2-9166, on EEG spectral content (Fig. 9J). While both drugs slightly decreased theta frequencies between 6 and 10 Hz, DZP increased all frequencies above 20 Hz, whereas LSP2-9166 significantly decreased the spectral content for frequencies above 14 Hz.

**Figure 9:**
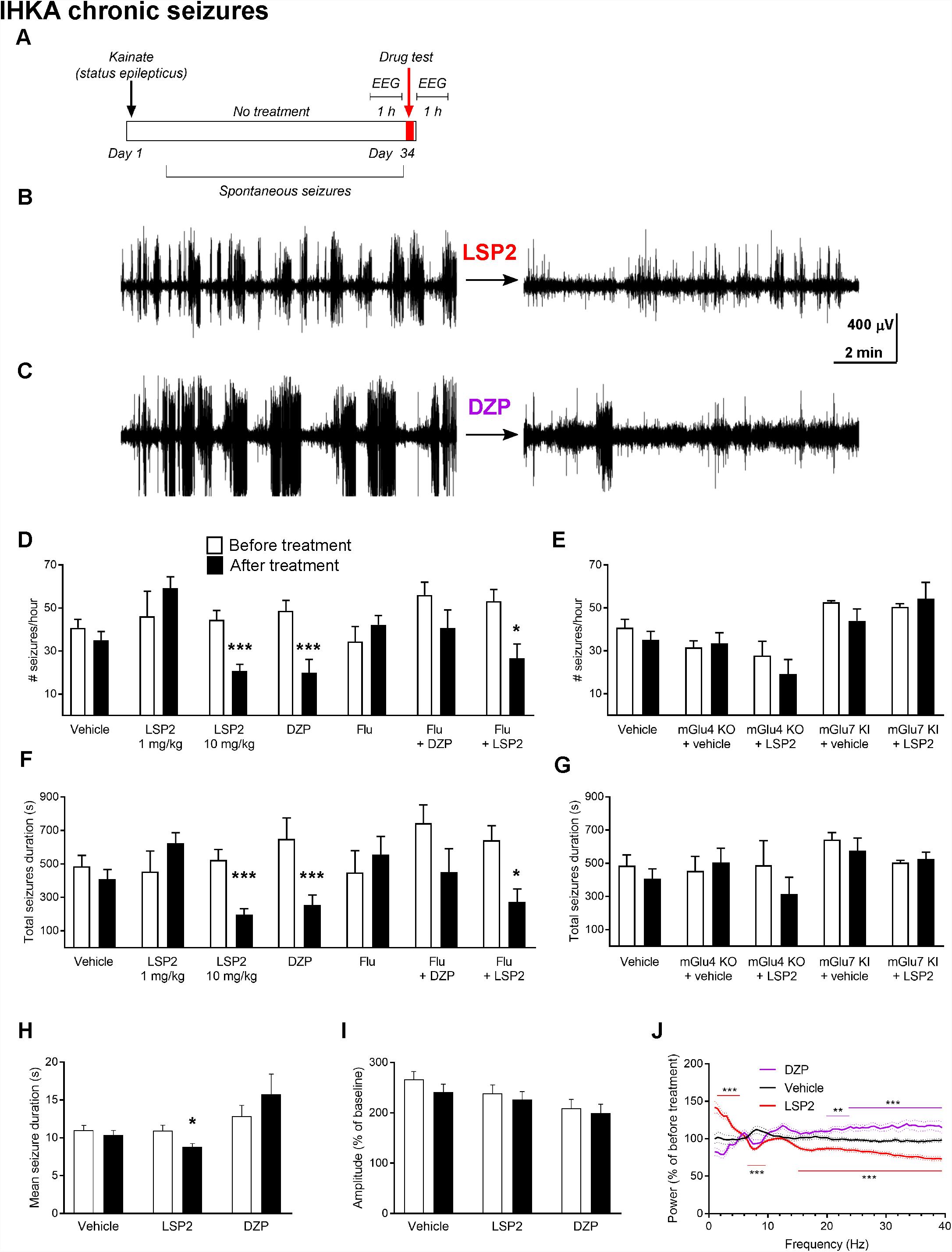
Anti-epileptic properties of LSP2-9166 on chronic IHKA seizures. (A) Experimental protocol. (B, C) Example of EEG traces representative of the acute LSP2 and DZP effect. (D-E) Number of seizures per hour depending on acute injection of compounds and genotypes. (F-G) Total seizures duration per hour depending on acute injection of compounds and genotypes. (H-I) Effect of acute compound injection on mean seizure duration and seizure amplitude. *, p< 0.05; ***, p<0.001, Mann-Whitney test. (J) Modifications of interictal power spectrum induced by either LSP2 or DZP. **, p< 0.01; ***, p<0.001, two-way ANOVA.

We evaluated the relative contribution of mGlu4 and mGlu7 receptors in blocking chronic hippocampal seizures by testing the LSP2-9166 on either mGlu4 KO or mGlu7 KI mice (Fig. 9E, G). Seizure number and duration showed a tendency to decrease only in mGlu4 KO mice, supporting the hypothesis that LSP2-9166 acts preferentially on mGlu7 receptors to counteract epileptic activity.

## DISCUSSION

### The effect of LSP2-9166 on epileptic progression and seizures

Looking for new targets to treat epilepsy is a long-standing challenge. We hypothesized that activation of mGlu7 has a potential in controlling epileptic progression and seizures. We took advantage of the development of a new orthosteric agonist acting on both mGlu4 and mGlu7, with an unmatched potency at mGlu7 receptor. To date AMN082 is the only mGlu7-selective agonist / positive allosteric agonist available. However, although AMN082 is orally active and brain-penetrating, it is not optimal for *in vivo* studies since it is rapidly metabolized and has off-target effects, notably on monoamine transporters (Sukoff Rizzo et al., 2011). LSP2-9166 has shown to be active in opiate addiction and alcohol consumption paradigms (Halassa et al., 2011; Lebourgeois et al., 2018).

We chose two complementary models of epileptogenesis with several advantages. Chemical kindling by repeated injection of PTZ allows the control of seizure occurrence: the behavioural response takes place within minutes after PTZ injection and no seizure is observed until the next administration of the convulsant. Nevertheless, the epileptogenesis process modifies the interictal brain activity, as shown by the modification of the basal EEG spectrum (Fig. 2E). Glial cell reactivity is also increased and creates a fertile ground for the progression of the network response. On the other hand, intra-hippocampal application of kainate provides a preclinically relevant mouse model of temporal lobe epilepsy, with focal hippocampal seizures, ipsilateral loss of CA1 pyramidal neurons, dentate gyrus dispersion, and inflammation as main hallmarks. Seizure progression is triggered by the initial status epilepticus (SE) and follows a maturation timeline over a few weeks, with full development of mature seizures at around 3 weeks post-SE. In the two models, we chose daily treatment with LSP2-9166 to face the continuous alterations taking place during epileptogenesis. The IHKA model being more severe, the compound was administered twice a day. In both cases, the treatment lasted 15 days, during which the majority of modifications are expressed. The aim was to define a therapeutic window to impact the epileptic network maturation. A period without treatment has then allowed to evaluate the stability of the effects, in order to differentiate between a temporary outcome and a long-term, disease-modifying effect.

Treatment with LSP2-9166 showed a remarkable effect on PTZ kindling, reducing to score 1 the response of the majority of animals. The effect was detectable since the beginning of the kindling process and was stable 30 days after interruption of the treatment, with no further compound administration. In addition, LSP2-9166 showed an AED-like effect on kindled seizures: a single dose of LSP2-9166 was able to reduce significantly the response of animals compared to day 15 of the kindling process. Interestingly, acute dosage of LSP2-9166 did not change the behavioural and EEG response to acute PTZ-induced seizures (Fig. 4A, B), whatever the PTZ dose. This results points at a real anti-epileptogenesis effect rather than a temporally limited block of PTZ antagonism on GABA-A receptors following each PTZ injection.

The fact that LSP2-9166 could not bring kindled animals below score 1 is *per se* remarkable: it could entail that the thalamocortical circuitry, which supports spike-and-wave discharges typical of the prostration status observed in score 1, is differentially affected by the compound compared to the rest of the epileptic network.

Overall, the animals treated with LSP2-9166 as soon as 6 h after SE showed a lower probability to develop seizures lasting longer than 15 s, as well as a shorter total seizure duration. Interictal periods are stretched and the number of seizures tends to be reduced compared to control animals. This tendency is constant over 15 days but the effect is less stable than for the PTZ kindling model. Indeed, 2 weeks after suspension of the treatment, the mean duration and total duration did not differ any longer from control animals. On the other hand, the effect on seizure-free time persisted beyond the end of the treatment. The limited effect of LSP2-9166 on the IHKA model might depend on an acute effect of the compound on seizures. These effects may be involved in an anti-epileptic - rather than anti-epileptogenic - effect. An alternative possibility is that the choice of day 15 post-SE to stop the treatment is not ideal, as it could be concomitant with a critical phase of network modification. As a fact, a loss of power in the theta band (4-12 Hz) was observed after day 15 post-SE in both treated and untreated animals (data not shown). Other therapeutic windows could be tested, to assess disease-modifying potential of the compound beyond the first epileptogenic events.

LSP2-9166 provided an AED-like effect on established, chronic IHKA seizures. Seizure number, mean duration and total duration, as well as the intrinsic frequency of seizures were significantly reduced over an hour post-administration, together with an increase in inter-seizure interval. On the other hand, peak amplitude was not affected by the treatment. A single dose of LSP2-9166 has an anti-epileptic effect as strong as diazepam (1 mg/kg). However, EEG power spectra analyzed during seizure-free intervals show a DZP-induced increase of frequencies above 20 Hz, while LSP2-9166 produces the opposite effect. The two drugs seem to have differential effects on brain activity, suggesting that the mechanisms at stake are not the same. In line with this observation, the benzodiazepine antagonist flumazenil counteracted DZP (Morag and Myslobodsky, 1982), but not LSP2-9166, in controlling chronic seizures.

### Effect on biomarkers

In the PTZ kindling model, inflammation is relative low (Kolosowska et al., 2014) and, as expected, we could not reliably detect different inflammatory mediators by RT-qPCR (including IL1beta, IL6, TNFalpha; data not shown). There were no changes in neurogenesis or cell proliferation. By contrast, there was a decrease in PDGFR-β mRNA, which could reflect a disruption of the blood-brain barrier (BBB) (Milesi et al., 2014). LSP2-9166-treated kindled animals showed a recovery of PDGFR-β mRNA levels. We observed neither cell loss nor neuronal reorganization as detectable by NeuN immunolabeling. However, modifications of astrocytic activation and microglia proliferation were detected in sub-regions of the hippocampus by immunohistochemistry. Changes in CA1 and CA3 microglia proliferation and CA3 and DG astrocyte activation were reversed by LSP2-9166. These results are in line with the neuroprotective effects suggested for mGlu7 activation in astrocytes (Jantas et al., 2018).

Compared to PTZ kindling, the IHKA model displays more sever alterations of hippocampal biomarkers, with significant differences between the ipsi- and contralateral injection side. Without treatment, as previously reported (Runtz et al., 2018), a striking increase in astrocytic activation and inflammatory response was observed after SE (Fig. 7). Accordingly, an increase of GFAP expression was detected in the ipsilateral side with an accentuation 28 days post-SE in the DG. Reactive astrocytes could be involved in guidance of granule cell migration (Bouilleret et al., 1999; Fahrner et al., 2007). An early increase followed by a later decrease in PDGFR-β mRNA levels, parallel the reported BBB disruption we previously reported in this model (Klement et al., 2018) and we observed that they were restored by LSP2-9166. A more detailed analysis of the agonist effect on the cerebrovascular integrity is ongoing. In the ipsilateral hippocampus an increase of mKi67 could reflect cell proliferation. By immunohistochemistry we observed both an increase in CA1 microglia proliferation as well as dispersion of DG, both attenuated by LSP2-9166. It should be noted that DG dispersion does not necessarily entails cell division but only cell migration along radial glia tracks (Fahrner et al., 2007). Altogether, our results suggest an important role of mGlu7 activation in neuro-glial reorganization, particularly in the DG.

### Repeated administration of LSP2-9166 is devoid of long-term side effects in naive animals

As part of this study, we evaluated the impact of LSP2-9166 administration on naive mice behaviour. Mice handling after injection suggested a slight change in the animal tonus, different from muscle tone loss or lethargy described after DZP injection or from the absence seizure-related prostration of mGlu7^AAA^ KI mice (Bertaso et al., 2008). In parallel, a cataleptic posture was observed when placing the animals’ forepaws on a rod. The effects reach a maximum 30 min after injection and decrease within 2 h, shadowing the plasmatic concentration of LSP2-9166. These effects could notably reflect a decrease in the level of vigilance, but a simple visual or auditory stimulation (movement or noise in the experimental room) was sufficient to resume the animal mobility. Similarly, testing animals on an active task as the rotarod showed no effect of LSP2-9166 treatment.

Repeated stimulation by the mGlu7 agonist could induce long-lasting functional modifications, as the receptor is implicated in synaptic plasticity in different brain areas including the hippocampus and the amygdala (Gee et al., 2014; Pelkey et al., 2008. However, a 15-day long daily systemic LSP2-9166 administration did not alter locomotor activity, coordination nor did it induce catalepsy. A slight decrease in the anxiety levels was detected in the elevate-plus maze test. A similar effect has been previously reported by either activation or inactivation of mGlu7 {Swanson, 2005 #857) and could provide an interesting avenue to treat an important comorbidity of epilepsy.

### Perspectives

Here we have shown that daily activation of mGlu7 acts on seizure maturation in both models of epileptogenesis. This protective effect deserves further investigation. Local injection of LSP2-9166 in different brain regions could allow the identification of areas susceptible to induce brain activity modifications similar to those observed upon systemic injection. In addition, if the brain/blood ratio of the compound parallels that of the corner stone compound, LSP1-2111 (Cajina et al., 2014), it will be interesting to assess the peripheral effects caused by the free fraction of the compound. For example, the mGlu7 receptor has been previously identified as an interesting new player in visceral pain or faecal water content (Julio-Pieper et al., 2010; Moloney et al., 2015).

Likewise, the contribution of mGlu4 receptors to the epileptogenesis process will need investigation in order to uncover the protective effect observed in the kindling model.

## MATERIAL AND METHODS

### Mice

All animal procedures were conducted in accordance with the European Communities Council Directive and approved by the French Ministry for Agriculture (2010/63/EU, authorization #746-2015072310069975). The generation and characterization of the mutant mGlu7 knock-in mice (mGlu7^AAA^ KI) and mGlu4 knock-out mice (mGlu4 KO) are described elsewhere (Pekhletski et al., 1996; Zhang et al., 2008).

### LSP2-9166 receptor activation

ThemGluorthostericagonistLSP2-9166,(2S)-2-amino-4-(((4-(carboxymethoxy)phenyl)(hydroxy)methyl)(hydroxy)phosphoryl)butanoic acid, was synthetized and characterized as described by (Acher et al., 2012). Compound activation of the different group III mGlu receptors (mGlu4, 6, 7, 8) was determined using transient transfection of each receptor in human embryonic kidney (HEK) 293 cells together with a chimeric G/Gq protein to allow measurement of calcium mobilization, as previously described (Tora et al., 2015).

### Plasma distribution of LSP2-9166

Mice were treated or not (0.9% NaCl solution) with LSP2-9166 at doses of 1 and 10 mg/kg (i.p.) and anesthetized with a mixture of ketamine (100 mg/kg) and xylazine (10 mg/kg) (i.p.) at different time points after drug injection (30 min, 1 h, 2 h, 6 h). Blood was then extracted directly from the heart by an intracardiac puncture and then transferred to 1.5 mL microcentrifuge tube containing 50 μl of a 5,000 UI/mL heparin sodium salt solution, centrifuged at 400 g for 15 min at 4°C and the resulting supernatant (plasma) was kept at −80°C until use.

The sample preparation and derivatization steps were previously optimized and the final conditions were as follows. 400 μL of −20°C pure methanol were added to 100 μL plasma. Thereafter they were vortex-mixed for several seconds and centrifuged at 13,000 g for 30 min at 4°C, and the resulting supernatant was separated and lyophilized using a centrifugal vacuum concentrator (Savant SpeedVac SVC-100). Samples were resuspended in 100 μL of pure water and 25 μL of the derivatization reagent were added by the autosampler equipped in the HPLC system. The derivatization reagent was prepared weekly by dissolving 25 mg of o-Phthaldialdehyde (OPA) in a mixture of 500 μL of ethanol, 5 ml of 100 μM borate buffer (pH 11.5) and 20 μL of 2-mercaptoethanol and was kept at 4°C and protected from light. Chromatographic separation was done on a reversed-phase Kromasil C18 column (250 mm× 4.6 mm, 5 μm particle size) purchased from Sigma-Aldrich. The Shimadzu Nexera system was equipped with dual pumps, an autosampler with a sample compartment set to 4°C and a column oven set to 45°C. Mobile phase A was 50 mM sodium phosphate pH 7.2. Mobile phase B was acetonitrile, pure methanol and water (2:2:1). Mobile phases were pumped at a flow rate of 0.800 mL/min using a gradient program that allowed for 2 min at 0% B followed by a 16-min step that raised eluent B to 60%. The total run time was 25 min. The injection volume was 40 μL. Fluorescence was monitored at excitation and emission wavelengths of 340 nm and 450 nm, respectively.

### Naive non-epileptic animals

Young adult male mice (6-7 weeks-old) were anesthetized with a mix containing ketamine (Imalgene 500, 100 mg/kg) and xylazine (Rompun 2%, 10 mg/kg) in PBS and placed in a stereotaxic frame using the David Kopf mouse adaptor. Mice were implanted a with a bipolar tungsten teflon-isolated torsade electrode positioned in the right dorsal hippocampus (AP = −2, ML = −1.5, DV = −2 mm from bregma (https://scalablebrainatlas.incf.org), as well as skull cortical electrodes on the frontoparietal bone and a reference electrode on the occipital bone. All electrodes were fixed on the skull with dental acrylic cement. After surgery, animals were individually housed and maintained in a 12 h light/dark cycle at 22 ± 2°C with food and water *ad libitum*. A first treated group received a daily intra-peritoneal injection of LSP2-9166 at 10 mg/kg in phosphate buffer solution, PBS, to guarantee proper pH and solubility during 15 days. A second treated group received a single intra-peritoneal injection of LSP2-9166 at 10 mg/kg in PBS. Control animals were injected with vehicle alone.

### Pentylenetetrazole (PTZ) kindling model

Young adult male mice (6-7 weeks-old) were anesthetized and prepared for surgery as described above. A female microconnector (Pinnacle Technology Inc., Lawrence, KS) was fixed by four extradural screws used as cortical electrodes, two on each parietal bone and then with dental acrylic cement.

Mice received a sub-convulsive dose of pentylenetetrazole (PTZ, 40 mg/kg i.p.) every other day for 15 days. The treated groups received a daily intra-peritoneal injection of LSP2-9166 at either 1 or 10 mg/kg in PBS. Control animals were injected with vehicle alone. Some of the animals were re-challenged with PTZ either 1 or 30 days after the last injection of mGlu7 agonist. A minimum of 60 min period of baseline EEG recording was allowed before injection of tested compounds. Electrical activity was recorded together with monitoring of animal behavior. The symptoms developing after injection were classified into different phases according to a modified Racines scale as described by (Weiergraber et al., 2006): phase 0 corresponds to normal exploration and posture; phase 1 corresponds to an “absence-like” non-convulsive state, with reduced motility and typical prostrated position of the animal; phase 2 shows partial clonus of head, vibrissae and/or forelimbs; phase 3 consists of a generalized clonus of the extremities, the tail, and sometimes vocalization; this phase can develop into typical “running fits”; phase 4 corresponds to a generalized tonic-clonic seizure with extension of the four paws; this phase is occasionally followed by death by respiratory failure.

For the observation of the effects of LSP2-9166 after acute injections, mice were prepared as described above and received one injection of either LSP2-9166 (10 mg/kg i.p.) or vehicle 15 min prior to a single injection of PTZ (30 to 60 mg/kg i.p.).

### Intra-hippocampal kainate (IHKA) injection model

Young adult male mice (6-7 weeks-old) were anesthetized with PBS-buffered solution containing chloral hydrate (Twele et al., 2016) and placed in a stereotaxic frame using the David Kopf mouse adaptor. A 10 µL microsyringe with stainless steel 33G beveled needle was filled with a 20 mM solution of kainic acid (KA; Sigma-Aldrich, France) in 0.9% sterile NaCl, and positioned in the right dorsal hippocampus (AP = −2, ML = −1.5, DV = −2 mm from bregma (https://scalablebrainatlas.incf.org). Mice were injected for 1 min with 50 nL (1 nmol) of the KA solution, using a micro-pump operating the micro-syringe. Sham mice were injected with 50 nL of 0.9% sterile NaCl under the same conditions. Right after intrahippocampal injection, mice were implanted at the same coordinates with a bipolar tungsten teflon-isolated torsade electrode, as well as skull cortical electrodes on the frontoparietal bone and a reference electrode on the occipital bone. All electrodes were fixed on the skull with dental acrylic cement. After recovery, the animals were kept under observation for 8–10 h after KA injection to visually determine their behavior during the KA-induced SE. The animals displayed mild asymmetric clonic movements of the forelimbs, clonic deviations of the head, rotations and/or prolonged periods of immobilization, and in some cases, bilateral clonic seizures of the forelimbs associated with rearing (Bouilleret et al., 1999). All mice displayed this characteristic behavioral pattern of SE after KA injection. Under these conditions the mortality was near to zero. A first treated group received a twice daily intra-peritoneal injection of LSP2-9166 at 10 mg/kg in PBS during 15 days post-SE. A second treated group received was used to study compound effects on chronic seizures at least 4 weeks post-SE. Control animals were injected with vehicle alone.

### EEG recording

EEG activity was recorded for all the animals mentioned above. Freely moving animals were put into individual Plexiglas boxes, and their microconnectors were plugged to an EEG preamplifier circuit and to the EEG amplifier (Pinnacle Technology Inc., Lawrence, KS, USA). Mice were first put to their test cage for 1 h, and were then regularly recorded for 2 to 24h. The electrical activity recorded by extradural electrodes was filtered at 40 Hz, sampled at 200 Hz and recorded by a computer equipped with Sirenia^®^ software (Pinnacle Technology Inc.). EEG recordings were performed together with video monitoring animal behavior.

To quantify the development of epileptogenesis, bipolar hippocampal and/or cortical derivations were used to record cerebral activities. In the IHKA model, the occurrence of hippocampal seizures, as described before (Riban et al., 2002), was analysed from day 5 post-SE. The threshold for detection of early epileptiform activity was set at 3-fold the baseline sighippnal. A seizure was defined as a paroxysmal activity lasting 5 seconds or more, and separated by at least 1 second from the next seizure, as previously reported (Duveau et al., 2016). Hippocampal seizures were detected using pClamp^®^ software feature of peaks detection and quantified to assess the number and cumulated duration per hour. Finally, the occurrence of epileptic activity was analyzed on the cortical derivation to assess whether hippocampal spikes or discharges propagated to the cerebral cortex. Power spectrum were obtained using Fast-Fourrier Transform (Neuroscore, DSI).

### Behavioural analysis

*Locomotor activity* was recorded using circular corridors (Imetronic, France). The circular corridor is composed of 4 infrared sensors (every quadrant) at ground level allowing recording of walking activities. Another detector is positioned at a height of 7.5 cm to follow mouse rearing activity. Horizontal activity is the number of quadrants that the mouse has crossed every 5 minutes while vertical activity corresponds to the number of rearing movements in a 5-minute period. Mice locomotion was recorded for a total of 60 min. *Motor coordination* was evaluated using a rotarod (Ugo Basile, Italy) to measure the latency to fall at either a steady speed of 6 rpm during 120 s or in the accelerating mode from 4 to 20 rpm over 120 s. *Catalepsy* was assessed by the latency to move from an unusual position after installing the mouse forepaws on a 4.5 cm height-rod. Activity in the *elevated-plus maze* was measured during 10 min as number of entries in the open/close arm and time spent in each type of arm.

### Immunohistochemistry

Mice were anesthetized using pentobarbital and transcardially perfused with 4% paraformaldehyde (PFA) followed by 24 h of post-fixation in 4% PFA. Thirty μm-thick slices were incubated in PBS-T (0.5 % Triton) with bovine serum albumin (3%) for 1 h and with primary antibodies (GFAP 1:300, IBA1 1:1000, NeuN 1:1000) for 12 h. Slices were then incubated for 2 h with secondary antibodies (AMCA anti-chicken 1:300, Alexa Fluor 488 anti-rabbit 1:1000, Cy3 anti-mouse) and DAPI. Acquisition was performed on an epifluorescence microscope. Laser power and detector sensitivity were kept constant for each staining to allow comparison between samples. At least 3 slices per animal were processed and images analysed using ImageJ software.

### Quantitative RT-PCR

Total RNAs were prepared using EZ-10 Spin Column Total RNA Miniprep Kit and treated with DNaseI according to the manufacturer’s instructions (Biobasic, USA), RNAs were quantified and their purity was checked at a Nanodrop (ThermoFisher, USA). RNAs were then reverse-transcribed into cDNAs using MMLV-RT enzyme and random hexamers (Life Technology, France). Quantitative PCR (qPCR) was performed in triplicate using SYBR Green Mix (Roche, France) on a Light Cycler LC480 Real-Time PCR system (Roche, France) in 384 well plates with 2 ng of equivalent RNA per point of qPCR. The level of expression of each gene was normalized to the mean of the expression levels of 3 housekeeping genes (*Hprt, Tbp* and *Gapdh*). These 3 reference genes were selected using the GeNorm procedure (Vandesompele et al. 2002). Quantitative-PCR primers are listed in Table 1.

**Table 1.**
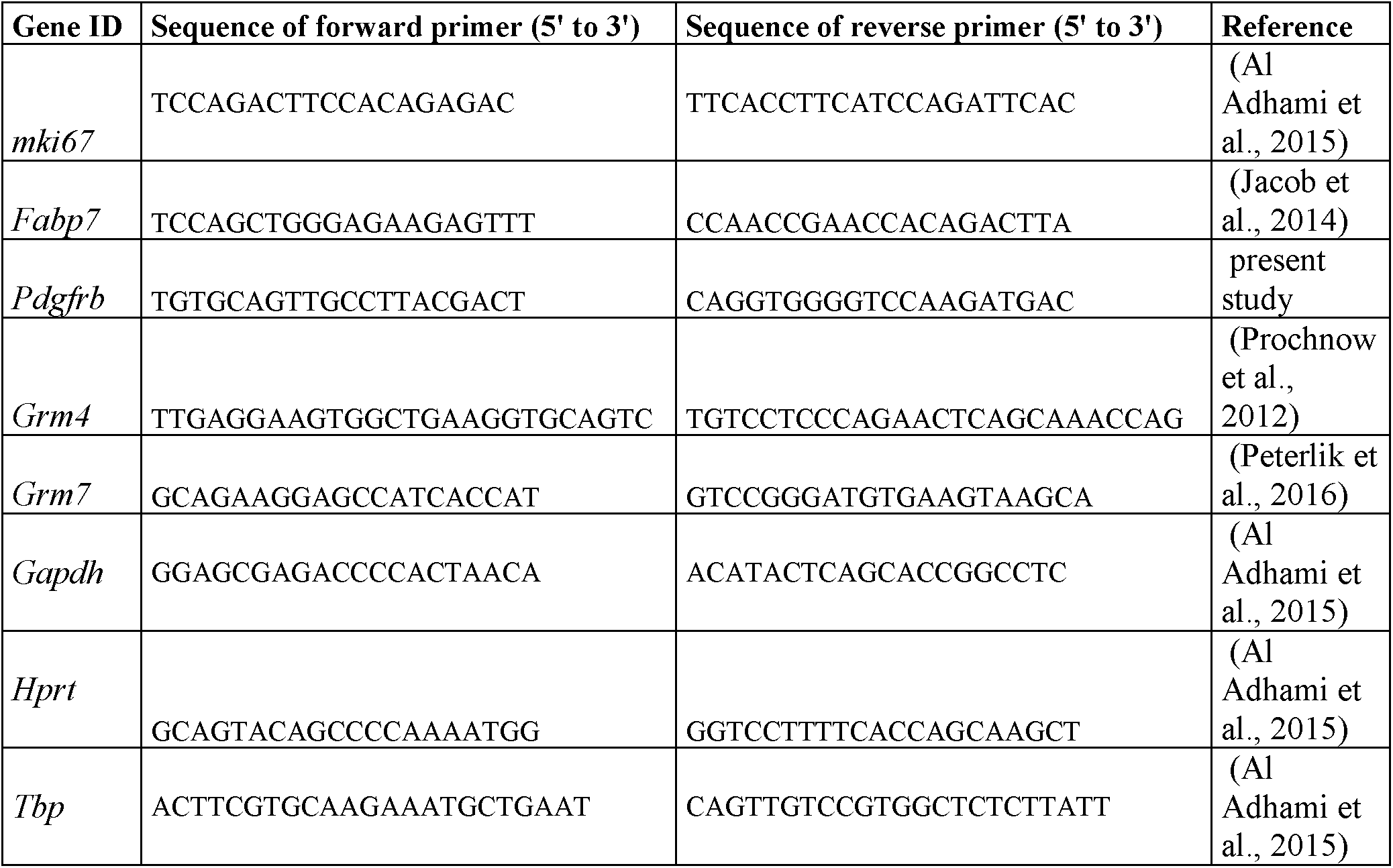
Primers for RT-QPCR

### Statistical analysis

Analyses were performed using pClamp 10, Neuroscore, GraphPad 7.05 and Microsoft Excel. Two-way ANOVA or Mann-Whitney’s tests were applied to determine significance with a 95% confidence interval. Power values for a priori sample size determination were obtained using G-Power open access software (http://gpower.hhu.de/).

## Acknowledgments

We are grateful to Isabelle Brabet and the Arpege platform for pharmacological tests, the iExplore animal facility at IGF, Antoine Depaulis (GIN, Grenoble, France) for help with the IHKA model, Cyril Goudet and Vanessa Pereira for providing mGlu4 KO mice. BG and FB designed the study, performed the experiments, analyzed the data, drafted the manuscript; PT, TB, AV, MPM, JV performed molecular analysis; JD designed and performed HPLC analysis; LF, NM, JP critically revised the data; CB, IM, FA designed the synthesis and synthesized LSP2-9166.

